# Identification of genes required for long term survival of *L. pneumophila* in water

**DOI:** 10.1101/2022.09.14.508055

**Authors:** Philipp Aurass, Seongok Kim, Victor Pinedo, Felipe Cava, Ralph R. Isberg

**Affiliations:** Department of Enteropathogenic Bacteria and Legionella, Robert Koch Institute, Wernigerode, Germany; Department of Molecular Biology and Microbiology, Tufts University School of Medicine, 150 Harrison Ave., Boston, MA. 02111; Laboratory for Molecular Infection Medicine Sweden, Department of Molecular Biology, Umeå Centre for Microbial Research, Umeå University, Umeå, Sweden

## Abstract

Long-term survival of *Legionella pneumophila* in aquatic environments is thought to be important for establishing an ecological niche necessary for epidemic outbreaks in humans. Eliminating bacterial colonization in plumbing systems is the primary strategy that depletes this reservoir and prevents disease. To uncover *L. pneumophila* determinants facilitating survival in water, a Tn-seq strategy was used to identify survival-defective mutants during 50-day starvation in tap water at 42°C. The mutants with most drastic survival defects carried insertions in electron transport chain genes, indicating that membrane energy charge and/or ATP synthesis requires the generation of a proton gradient by the respiratory chain to maintain survival in the presence of water stress. In addition, periplasmically-localized proteins that are known (EnhC) or hypothesized (*lpg1697*) to stabilize the cell wall against turnover were essential for water survival. To test that the identified mutations disrupted water survival, candidate genes were knocked down by CRISPRi. The vast majority of knockdown strains with verified transcript depletion showed remarkably low viability after 50-day incubations. To demonstrate that maintenance of cell wall integrity was an important survival determinant, a deletion mutation in *lpg1697*, in a gene encoding a predicted L,D-transpeptidase domain, was analyzed. The loss of this gene resulted in increased osmolar sensitivity and carbenicillin hypersensitivity relative to the WT, as predicted for loss of an L,D-transpeptidase. These results indicate that the *L. pneumophila* envelope has been evolutionarily selected to allow survival under conditions in which the bacteria are subjected to long-term exposure to starvation and low osmolar conditions.

**Importance:** Water is the primary vector for transmission of *L. pneumophila* to humans and the pathogen is adapted to persist in this environment for extended periods of time. Preventing survival of *L. pneumophila* in water is therefore critical for prevention of Legionnaire’s disease. We analyzed dense transposon mutation pools for strains with severe survival defects during a 50-day water incubation at 42°C. By tracking the associated transposon insertion sites in the genome, we defined a distinct essential gene set for water survival and demonstrate that a predicted peptidoglycan crosslinking enzyme, *lpg1697*, and components of the electron transport chain are required to ensure survival of the pathogen. Our results indicate that select characteristics of the cell wall and components of the respiratory chain of *L. pneumophila* are primary evolutionary targets being shaped to promote its survival in water.

## Introduction

*Legionella pneumophila* (Lpn) is a water-borne pathogen that is the causative agent of Legionnaire’s disease, which results in pneumonia associated with high levels of lethality (1). The bacterium and related species are widespread in natural and man-made water systems, it persists planktonically, colonizes biofilms and infects permissive protozoa (2, 3). The colonization of municipal water systems is of great significance (4), as it is the leading cause of drinking water-related Legionnaire’s disease outbreaks in the US (5). This facilitates infection via the main route of transmission to humans, which is inhalation of contaminated aerosols shed from poorly managed water systems. Once contaminated, eradication of *Lpn* from water supplies has proven challenging, so much of public health containment has been devoted to engineering water systems that prevent colonization of the pathogen. In the past two decades, there has been consistent increase in the number diagnosed Legionnaire’s disease cases (6). The estimated direct healthcare cost of Legionnaires’ disease due to hospitalization and ED visits in the United States was estimated over USD 400 million in 2014 (1).

As a waterborne pathogen, Lpn is equipped to survive, persist, and when associated with biofilms, grow in drinking water (4). It has the documented ability to persist in water for months or even years without losing its ability to regrow and to infect host cells (7–10). As Legionnaire’s disease is highly seasonal in nature, this indicates that the bacterium may persist for very long periods of time in water supplies prior to an epidemic outbreak of disease. The expanding burden of waterborne Legionellosis illustrates the need to better understand how *Legionellae* survive in water and in protozoa in order to identify points of vulnerability, and to design remediation strategies.

Previous work highlighted the importance of the stringent response and the alternative sigma factor RpoS for starvation survival in water. Key factors orchestrating its immediate adaptation to nutrient limitation include the LetA/S-rsmYZ-CsrA regulatory cascade (11–14). The CsrR protein promotes long term survival by a yet unknown mechanism (15). Other water stress survival-promoting factors include a putative metal transporter LasM and a putative transcriptional regulator Lpg2524 (16, 17). Transcriptomic or proteomic screens identified Lpn genes upregulated under prolonged exposure to water (“water stress”) or proteins particularly abundant under these conditions (18, 19). These studies were designed to positively select for the (over)expressed subgenome under water stress conditions. Weakly transcribed or repressed genes with importance during persistence or for regrowth might escape these screening approaches.

Most of the effort in understanding the physiology of Lpn has been devoted to identifying and determining the mechanism of action of proteins necessary for intracellular growth of the bacterium within environmental amoebae and target macrophages, which form the replicative niche during pneumonic disease in humans (20, 21). In contrast, there has been little progress in identifying proteins necessary for survival in tap water, partly because systematic analysis requires extensive incubation under starvation conditions. To overcome this block, we identified the entire spectrum of proteins involved in supporting survival after long-term incubation in water stress conditions. To this end, we used transposon insertion sequencing (Tn-seq) to identify mutants with low fitness in the presence of starvation in sterilized tap water (22).

In this report, we identify proteins not previously known to support survival during water starvation (10, 14, 23), including a protein we show is critical for infection of an amoebal host as well as mammalian macrophages. Our study expands the current knowledge on persistence factors of Lpn and reveals new points of vulnerability for the organism during persistence in water.

## Results

### Identification of L. pneumophila genes required for survival during iron starvation

To identify candidate genes that are specifically essential for long-term survival of *L. pneumophila* in tap water but viable during growth in bacteriological medium, we employed mutagenesis with the mariner transposon Himar-1, which inserts randomly in chromosomal TA sites (24). From the pools generated, we used Tn-seq, in which massive parallel sequencing of transposon-chromosome junctions was performed to detect the locations of insertions before and after water starvation. We generated three independent mutant libraries with 64,000, 75,000, and 87,000 distinguishable members. This is equivalent to 26%, 30%, and 36% of the 248,087 theoretically possible integration sites, both within and outside annotated open reading frames (ORFs) (25). These three Tn-seq library datasets were then analyzed as a combined superpool representing a total of 54,491mutants with distinct transposon insertions in ORFs, which represents 1/3 of the 166,297 theoretically possible ORF AT sites (NCBI SRA PRJNA816157).

The mutant libraries were subjected to water stress at 42°, taking advantage of the fact that elevated temperatures are known to accelerate viability decay in water, shortening the time required to analyze the datasets (9, 26) (Fig. 1). To this end, 300,000 colony forming units (cfu) for each library were incubated on solid bacteriological medium and harvested either directly after tap water inoculation (t1) or after 50 days water exposure (t50). The CFU/ml in water of each library dropped by 60% during the 50-day incubation (from 1×10^8^ to 4×10^7^ CFU / ml). Genes specifically necessary for high fitness under starvation conditions were discovered by using the Bayesian/Gumbel method embedded in the TRANSIT software package (22), identifying mutants that were poorly recovered at 50 days relative to their abundance on initial inoculation into water (Fig. 1). This approach identifies open reading frames with long stretches showing much lower insertion density than observed prior to water incubation. The probability of such insertion gaps occurring by chance is then calculated by a Bayesian analysis of the Gumbel distribution, and essential (poor representation relative to initiation of incubation), non-essential, and “uncertain” or equivocal phenotypes are called (22).

**Fig. 1.**
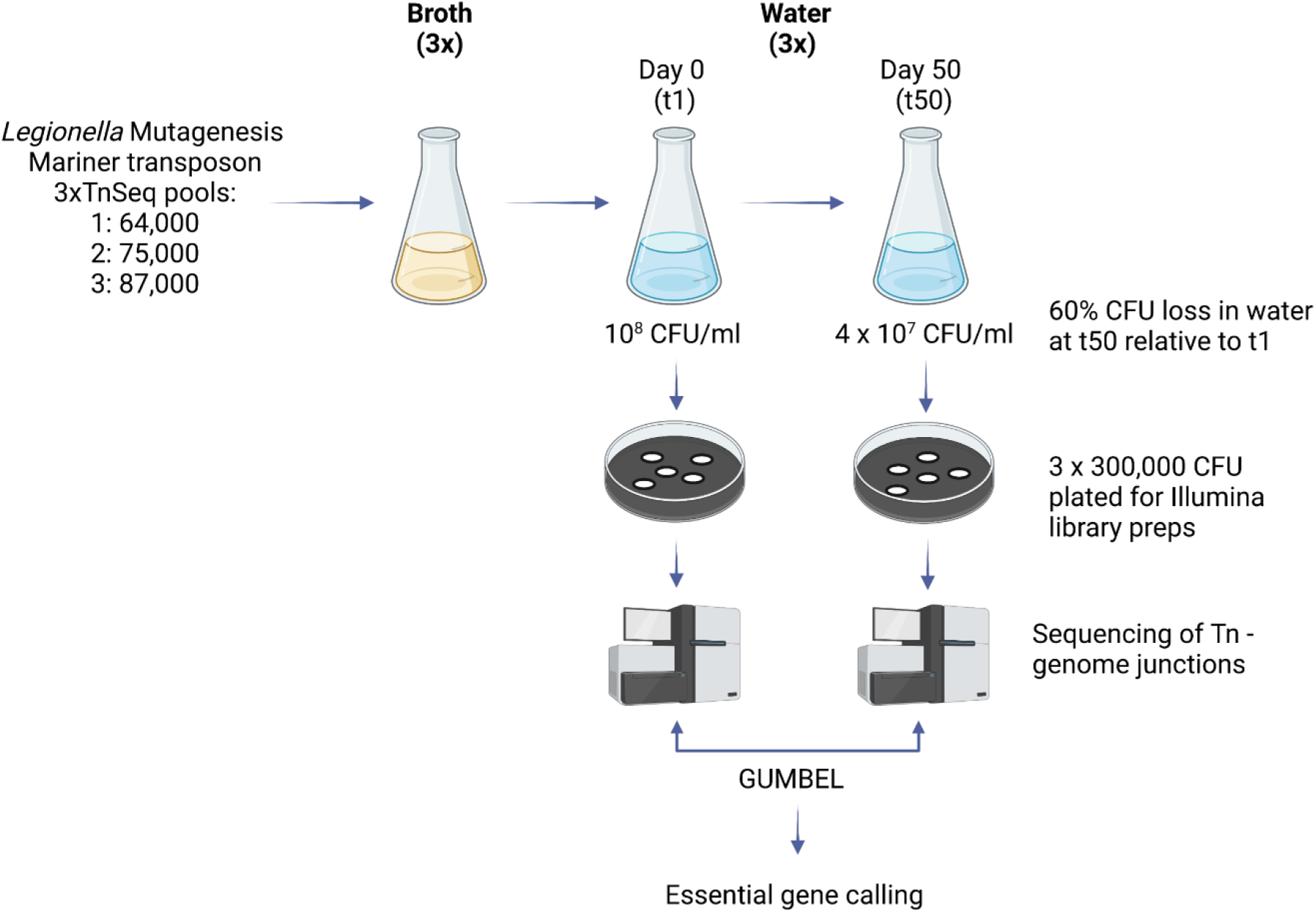
Tn-seq screening strategy for identification of *Legionella pneumophila* genes required for long term survival in water. Three separate Tn-seq pools with 64,000 to 87,000 individual bacterial members were generated using the mariner transposon Himar-1. The individual pools were expanded to postexpoential growth phase in BYEα broth (OD600 3.8) and then inoculated to sterilized tap water at an initial density of 10 CFU / ml. After 50 days of incubation at 42°C, the CFU in water had dropped by 60 % relative to t1. At t1 and t50, 300,000 CFU per library were plated and harvested for gDNA isolation and Illumina library preparation after 2 days of growth. Libraries were sequenced using Illumina HiSeq and frequencies of transposon insertions per gene were determined at t1 and t50. The frequencies of insertions were subsequently statistically analyzed with GUMBEL to call essential genes at the t1 and t50 (22). Figure was created with BioRender.com.

Based on the analysis of the Tn-seq superpool, out of a total of 2943 annotated *L. pneumophila* Philadelphia 1 ORFs (NC002942), 583 genes were called essential (E) after growth in bacteriological media (22) (Fig. 2A, Tab. S1). When this pool was subjected to water stress for 50 days, the essential set included 32 additional genes (Fig. 2B, red). We also included genes with a more than 3-fold drop in abundance at t50 relative to t1 in our analysis to account for genes with severely reduced fitness after the treatment, but did not reach the threshold of significance at the employed library saturation level (Fig. 2B, green datapoints, Tab. S1). The combined gene sets resulted in a list of 45 genes (Tab. 1), which we defined as the candidate gene set required for survival after prolonged starvation of *L. pneumophila* in water. The candidate genes were ordered according to their known or predicted functions and overrepresented in functions associated with the bacterial envelope (Table 1). These included factors involved in respiration (12 genes), membrane transport (8), hypothetical proteins (6), regulation (6), “virulence” (3), DNA replication, repair, and recombination (3), intermediary metabolism (5), stress (1), and protein processing (1) necessary for high viability under water stress conditions. Of particular note is the reidentification of *rpoS* sigma factor and the *letA/S* regulatory system as being essential under water starvation conditions. As these three genes had been previously discovered as controlling survival under water starvation, this argues that our selection procedure reproduces previously established starvation conditions in the literature (14, 23).

**Fig. 2.**
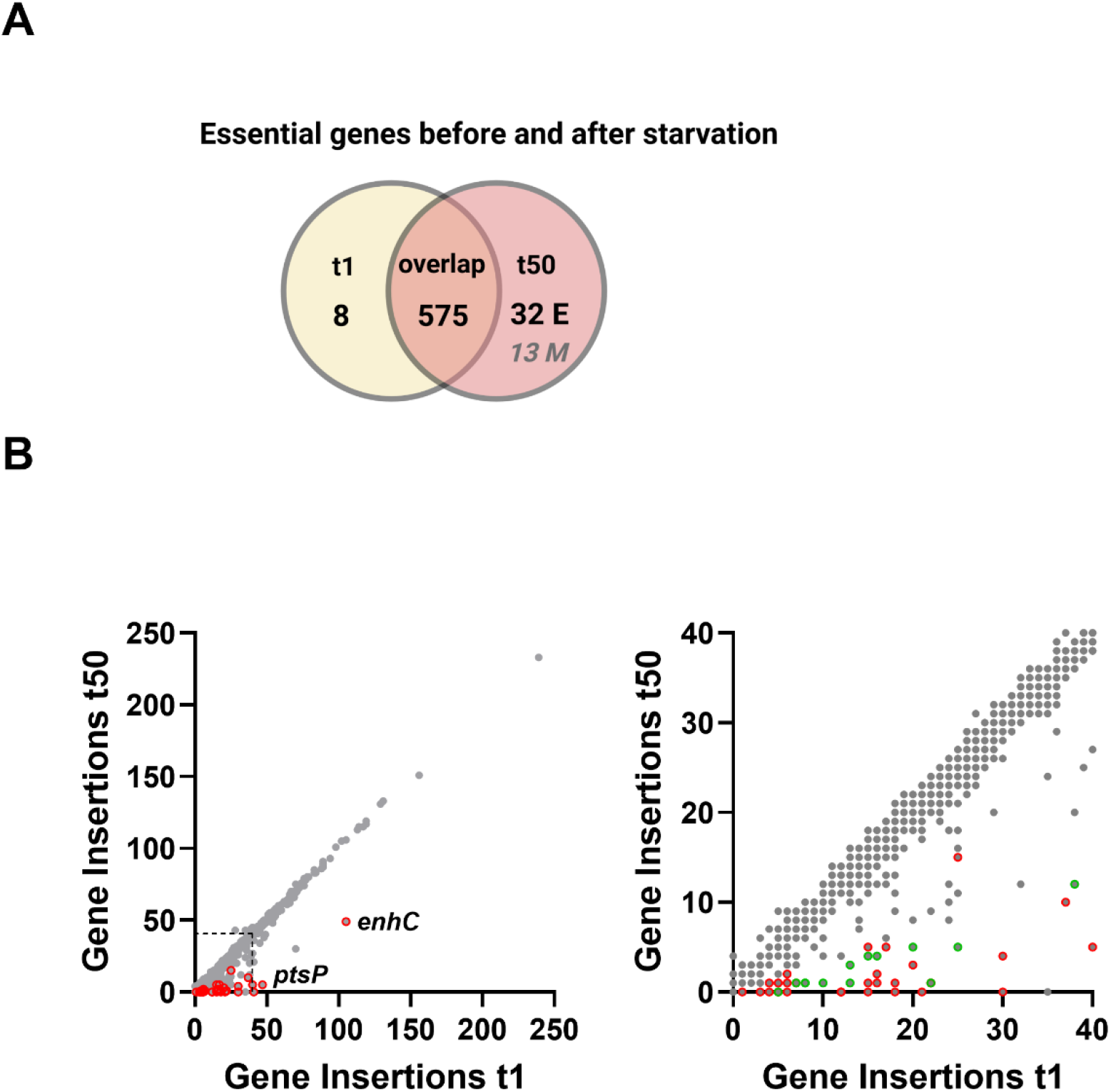
Strategy for identification of 45 genes critical for survival during long term water stress. Identification of essential genes after 50 days starvation in water at 42°C compared to bacteriological medium followed by short water exposure. A) GUMBEL analysis identified 575 genes essential [E] both after immediate addition into water (t1, yellow circle) or incubating for 50 days at 42°C (t2, red circle; overlap). By manual selection [M], we identified 13 additional genes presumably important for starvation survival in t50 that where below the threshold of significance in GUMBEL (italic number, M). B) Comparison of insertion abundance/gene for bacteria at t50 vs t1. Left diagram shows all genes (gray) with the 32 essential genes called by GUMBEL encircled in red. Right diagram shows the 0 to 40 insertions/gene region only (boxed in left side). Red circles indicate essential genes under water starvation conditions called by GUMBEL. Green circles indicate additional genes presumably important for starvation survival based on low abundance after 50-day water exposure. Members of the additional gene set had a minimum of 5 insertions at t1. After starvation, the abundance of insertions occurring in members of this latter group was reduced by more than 3-fold (13 genes). Fig. 2A was created with BioRender.com.

**Tab. 1.**
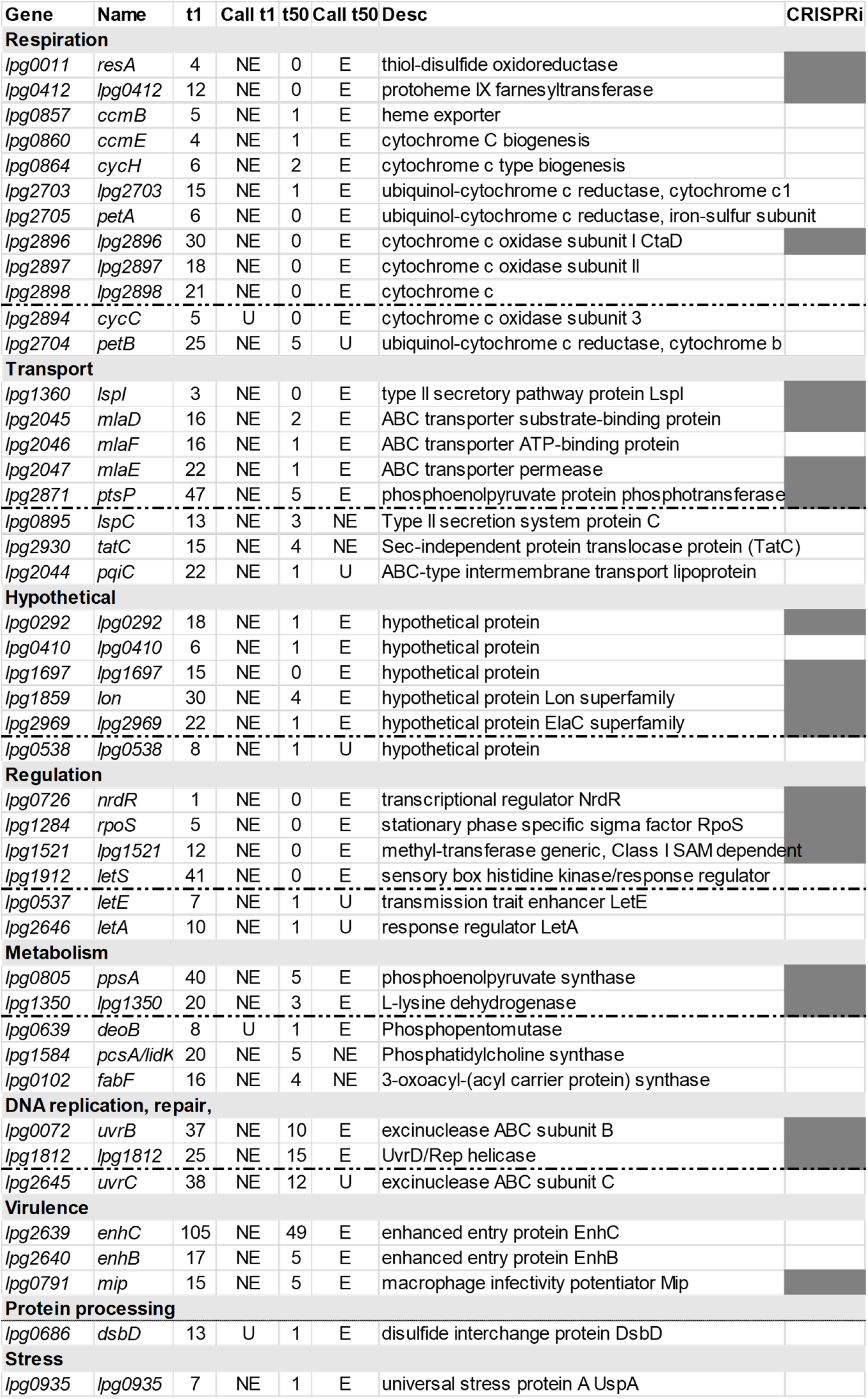
Candidate gene set required for survival after prolonged starvation in water. t1/t50=number of insertions per gene observed at timepoint. Call t1/t50: E=essential, NE=nonessential, U=uncertain. CRISPRi: grey shaded tiles indicate genes chosen for validation by knock down. Manually identified genes are separated by dashed line at the end of each section (Fig. 2b). Further details listed in Tab. S1.

### Efficient knockdown of gene transcription with CRISPR interference

We next directed our analyses to individual members of the candidate list of water stress survival genes, focusing on proteins not previously noted as important for survival in water starvation conditions (Table 1), to validate the predictions made by our Tn-seq analysis. For this purpose, we took advantage of the recently established CRISPR interference system for *L. pneumophila* (27). This system utilizes an anhydrotetracycline (ATC)-inducible *dcas9* gene inserted in the nonfunctional chromosomal *thyA* locus of *L pneumophila* strain Lp02, and crRNA-as well as tracrRNA-encoding sequences on a plasmid (27, 28).

Previous work has demonstrated that CRISPRi technology allows efficient gene knockdowns in *L. pneumophila* during growth in either broth or phagocytic cells (27). Its applicability to prolonged water starvation conditions has not been previously established. Therefore, we tested a knockdown in a gene encoding an Icm/Dot component, which has no effect on long-term survival in water (NCBI SRA PRJNA816157, Tab. S1) but blocks intracellular growth, to determine the effectiveness of this strategy. This allowed us to determine if long-term water incubation altered the efficiency of ATC-activated CRISPRi depletion. To this end, a *crdotO* strain, which showed a 30-fold knockdown efficiency relative to control Lp02 when assayed after growth to postexponential (PE) phase, was tested for its ability to block growth within *Acanthamoeba castellanii* after water starvation (Fig. 3A). To minimize the adverse effects on bacterial viability but maximize the timescale we performed this experiment at ambient temperature (22°C), starving for either 35 or 70 days (Fig. 3).

**Fig. 3.**
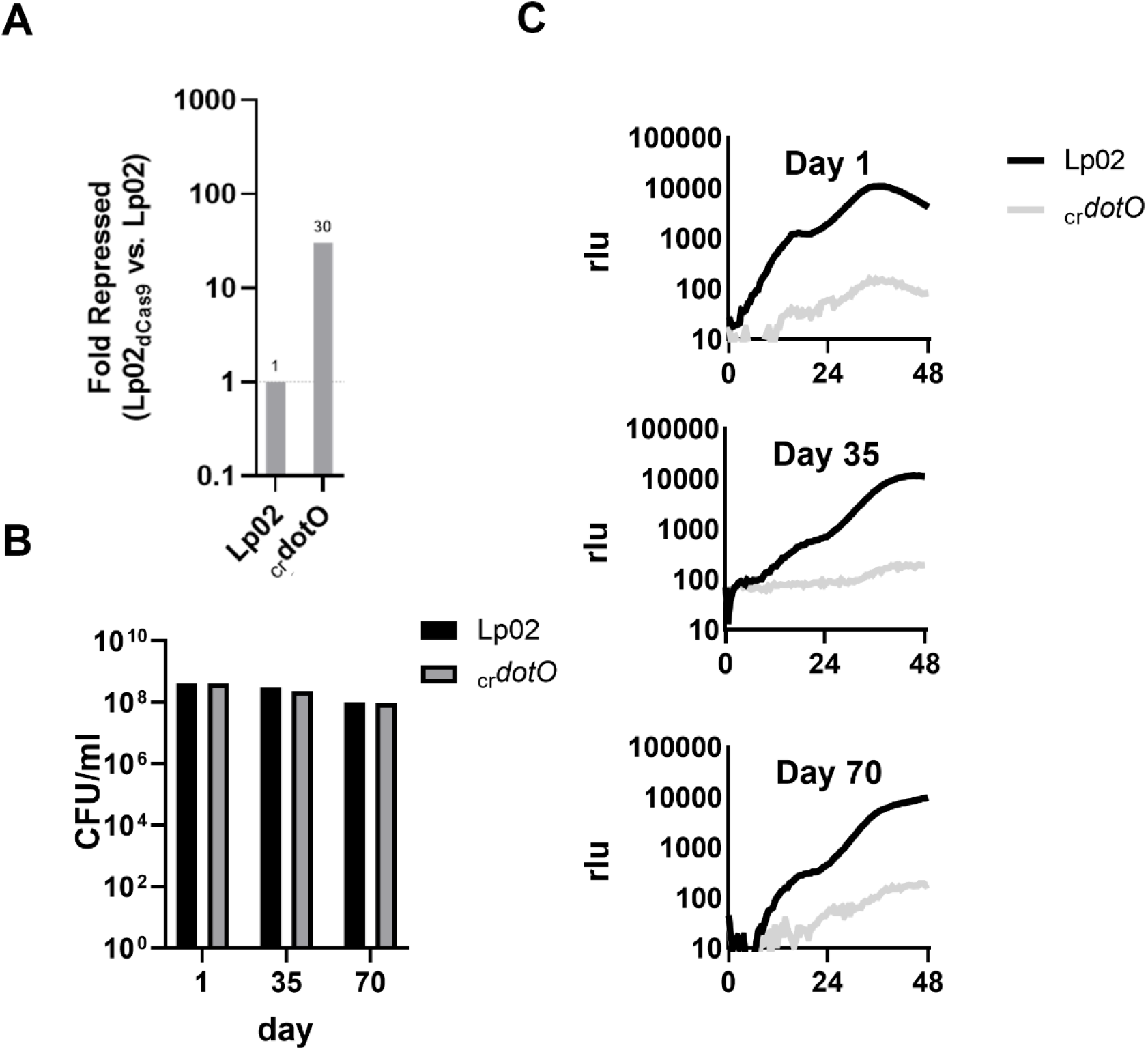
CRISPRi strategy results in efficient depletion and allows blockade of intracellular replication under starvation conditions. A) Reduction of steady state transcript levels by CRISPRi after growth in broth and inoculation in water (t1). Lp02dCas9Lux(_cr_*dotO*) was grown in presence of ATC inducer in broth, inoculated in water, and transcription levels were compared to Lp02 (WT) by qRT-PCR. B) High viability of Lp02Lux and _cr_*dotO* strains during extensive water exposure. Viability was measured by spot plating CFU/ml on BCYEα agar. C) CRISPRi interference of *dotO* blocks intracellular growth. Bacteria were inoculated in tap water for noted times, then used to challenge *A. castellanii* over 72 hours. Intracellular growth was measured by luminescence. Multiplicity of infection used (MOI) = 0.5. rlu=relative light units.

The effects of the *dotO* CRISPRi knockdown were followed phenotypically, as gene expression analysis in starved bacteria is made difficult by the largescale drop in transcription and changes in gene expression dynamics that occur in these conditions relative to broth-grown bacteria (29). After growth in broth to postexponential phase, there was a 30-fold drop in steady state levels of *dotO* transcript in the presence of *dotO* crRNA (Fig. 3A). As predicted by the Tn-seq dataset, this caused no reduction in viability relative to the WT at either 35 or 70 days of tap water starvation (Fig. 3B). In contrast, *dotO* CRISPRi knockdown resulted in a total block in growth within *A. castellanii* of Lpn harboring a luciferase reporter, with the level of defect indistinguishable for bacteria undergoing long-term water starvation (35 or 70 days) or for bacteria assayed shortly after removal from broth and washing in water (Fig. 3C). These data demonstrate that we could separate water survival from defective intracellular growth and that CRISPRi knockdown persists over long-term water incubation, allowing phenotypic assays to be used *in lieu* of a direct measure of transcription after starvation. We conclude that the CRISPRi system (27), in addition to efficient gene knockdown in broth-grown Lpn, can be used to evaluate starvation survival and infection dynamics after long-term water stress.

### Demonstration that candidates are required for survival under water stress conditions

Having demonstrated the combined applicability of CRISPRi and the bacterial luciferase operon as an indicator of intracellular survival, we next created single crRNA-encoding plasmids for specific candidate water stress survival genes as well as a nontargeting sequence derived from the mRFP orf. These were then introduced into the WT strain Lp02dCas9Lux (30) (Table 1, Supplementary Table S2). This strain has the bacterial luciferase operon *luxCDABE* downstream of the chromosomal *PahpC* in the *thyA::dCas9* background (31, 32) allowing the downstream analysis of metabolic activity of CRISPRi-depletion strains. The luciferase system uses cofactors, such as NADPH, ATP, FMNH2 and acetyl-CoA, allowing it serve as an intrinsic proxy for metabolic activity (31, 33). Observed loss of luciferase activity is likely due to loss of metabolic activity, so absence of activity after metabolic stimulation by laboratory growth media or after coculture with permissive host cells, is a signal that the bacteria are presumably dead.

As our Tn-seq experiment relied on the ability of a mutant to compete against a large population, we wanted to test if the candidates simply showed poor competition but otherwise were fit when incubated in isolation during long-term water starvation. To address this point, individual crRNA derivatives were grown in broth in the presence of ATC and then subjected to water starvation as described for the *dotO* depletion. Analysis was directed toward 18 candidates hypothesized to be involved in water stress survival, picking representative members of each functional group, as well as the known water stress survival gene *rpoS* as a control. These were chosen to avoid independent testing of multiple genes belonging to the same functional group, such as the respiratory chain genes (Tab. 1) (23), and already characterized stress survival genes such as *letS* and *uspA*.

Bacteria were incubated in tap water at 42°C, and at various times over a 50-day period, aliquots were serially diluted onto solid bacteriological medium to quantitate viable bacteria, comparing CFU to the nontargeting control (Fig. 4A). 18 days of water incubation resulted in a number of the candidates showing a loss of viability relative to the nontargeting control (Fig. 4A, charcoal triangles). Most striking was crRNA directed against *lpg2969*, which showed no detectable CFU after 18 days incubation (Fig. 4A; 10^-7^ CFU relative to nontargeting control). 32 days in water starvation conditions resulted in decreased viability of 67% of the strains that were tested, with depletions of *mip, rpoS, ccmH*, and *ctaD* also showing no viability at this timepoint, with extremely poor viability of the *lon, lpg1697*, and *mlaD* depletions. At 50 days in water, depletions of the additional candidate genes *lpg0011, mlaD, lon, lpg1521, ptsP, mlaE, lpg1697, lpg1350* showed no viability on plates. As the majority (68 %) of the depletion derivates resulted in absence of viability in water, our selection procedure identified mutants that are involved in tolerating water starvation conditions even in the absence of competition with the whole populations (Table 2).

**Fig. 4.**
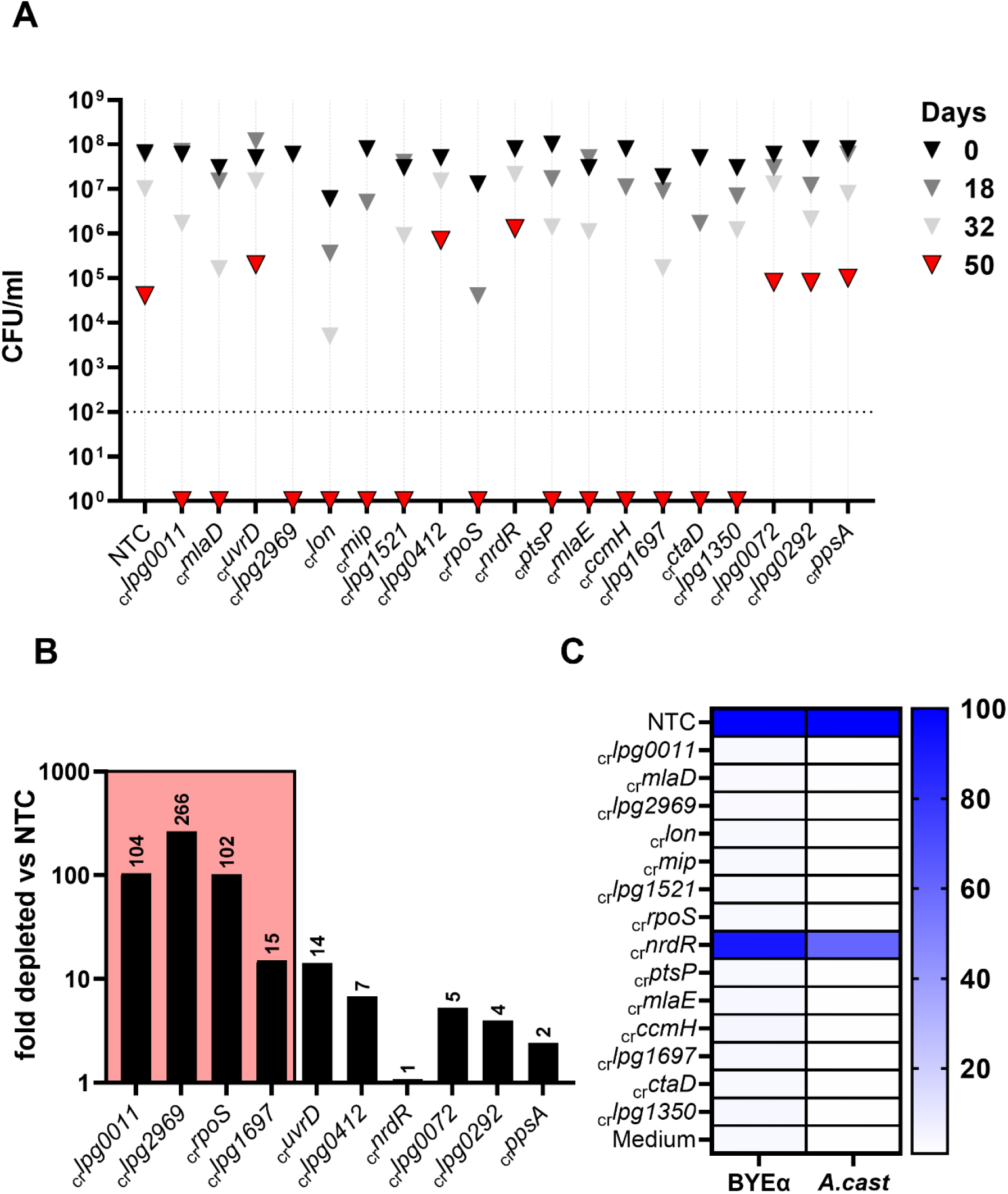
CRISPRi knockdown results in loss of viability of water stress survival-defective candidates after long-term water exposure. A) CFU/ ml of 19 candidate CRISPRi knockdown strains identified as conditionally essential by Tn-seq after extensive water stress. Dotted line=detection limit. Triangles: black=0 d., dark gray=18 d., gray=32 d., red=50 d. Each strain represents a single clonal water culture. Data show one out of two independent experiments with similar results. B) Efficiency of CRISPRi depletion correlates with stress survival phenotype. Analysis of knockdown efficiency in select knockdown strains relative to the non-targeting control. Strains were chosen based on culture phenotype, including four culture-undetectable strains after water incubation (coloured box), and the six strains that still formed colonies after 50 days water incubation. C) Culture in *A. castellani* does not rescue CRISPRi candidates that are not viable in bacteriological medium. Heat map of RLU expressed as area under curve after 24h (BYEa) or 72h (*A. cast*) of growth in BYEa or during coculture in *A. castellanii* of culture negative strains as well as NTC and _cr_*nrdR* culture-viable controls.

**Tab. 2.** List of starvation survival essential genes identified by Tnseq and confirmed by CRISPRi knockdown. *NCBI CDD database, **known or inferred from CDD

| gene | name | CD* [name / accession] | localization** | function** | PMID |
| --- | --- | --- | --- | --- | --- |
| *lpg0011* | thiol-disulfide oxidoreductase | TlpA_like_family | periplasm | energy metabolism |  |
| *lpg0791 / mip* | macrophage infectivity potentiator | FkpA / COG0545 | membrane | pathogen | 1594630 |
| *lpg0864 / ccmH* | cytochrome c type biogenesis protein CcmH | NrfG super family | periplasm | energy metabolism |  |
| *lpg1284 / rpoS* | stationary phase specific sigma factor | rpoS_proteo / TIGR02394 | cytosolic | regulation | 25416763 |
| *lpg1350* | L-lysine dehydrogenase | Lys9 super family | cytosolic | lysine metabolism |  |
| *lpg1521* | methyl-transferase | AdoMet_MTases super family | cytosolic | regulation / modification |  |
| *lpg1697* | hypothetical protein | YkuD_2 / pfam13645 | periplasm | peptidoglycan modeling |  |
| *lpg1859 / lon* | ATP-dependent protease La | Lon / COG0466 | cytosolic | protein homeostasis | 17891379 |
| *lpg2045 / mlaD* | ABC transporter substrate-binding protein | MlaD / COG1463 | inner membrane | membrane homeostasis |  |
| *lpg2047 / mlaE* | ABC transporter permease | MlaE / STAS superfamily | trans-membrane | membrane homeostasis |  |
| *lpg2871 / ptsP* | phosphoenolpyruvate protein phosphotransferase | PtsP super family / cl34645 | inner membrane | energy metabolism |  |
| *lpg2896 / ctaD* | cytochrome c oxidase subunit I | Cyt_c_Oxidase_I / cd01663 | inner membrane | energy metabolism |  |
| *lpg2969* | hypothetical protein | ElaC / COG1234 | cytosolic | translation |  |

6 of the 19 candidates chosen for validation by CRISPRi resulted in approximately the same fraction of survivors after 50 days starvation in water as the nontargeting control. To test if absence of phenotypes in these strains could be due to insufficient target depletion by dCas9, we performed qRT-PCR with RNA isolated from the culture aliquots obtained at the onset of water cultures after growth in broth. In addition, we chose four confirmed candidate genes and the NTC for reference (Fig. 4B). Transcript depletion of the confirmed genes was 104-fold (*lpg0011*), 266-fold (*lpg2969*), 104-fold (*rpoS*), and 15-fold (*lpg1697*). In contrast, the depletion levels of the targeted transcripts from the strains showing no apparent survival defect was lower, ranging from 14-fold (_cr_*uvrD*) to no effect (_cr_*nrdR*) (Fig. 4B). These data are consistent with inefficient transcript depletion being the cause of the absence of detectable phenotypes in these strains.

We next determined if the strains showing low viability on solid medium were metabolically inactive based on loss of luciferase activity and restriction by *A. castellani*. After 50 days of starvation, culture aliquots of the 13 strains that showed no viability on solid medium at this time point were metabolically stimulated and cocultured with *A. castellanii* in an attempt to resuscitate unculturable *L. pneumophila* (Fig 3C). The control strains (NTC and the *_cr_nrdR* harboring strains) readily responded to metabolic stimulation with increased luciferase activity, and were capable of intracellular growth (Fig. 3C). In contrast, there was no evidence of luciferase activity in the strains tested and none of these were resuscitated by coculture with amoebae. Since no signs of metabolic stimulation through nutrient addition or coculture were evident in the knockdown strains (Fig. 3C), we conclude that knockdown of this particular set of genes resulted in no evidence of resuscitation under any condition and their loss caused bacterial cell death.

We conclude that, as predicted by the results of the original selection, overrepresented processes such as maintenance of bacterial envelope homeostasis (*mip, lpg1697*, and *mlaDEF*), electron transport (*ctaD*) and cytoplasmic protein turnover (*lon*) are critical components in supporting survival in the face of water stress (Table 2). The consequence of loss of these proteins, which are not essential during growth in bacteriological medium, is bacterial death after 50 days incubation in tap water.

### Lpg1697 *carrying a YkuD domain is required for survival in presence of water stress*

Lpg1697 encodes a predicted 225 aa polypeptide carrying a predicted L,D-transpeptidase domain (LDT; YkuD_2), and a predicted N-terminal signal peptide (SignalP-5.0). Previous studies showed the peptidoglycan layer of Lpn is highly crosslinked (34) and as many as 10 LDTs have been predicted to be encoded by the genome of *L. pneumophila* (35) (Fig S1). Unlike characterized LDTs of other genera, Lpg1697 lacks a predicted LysM peptidoglycan-binding domain, questioning its functionality as a murein modifying enzyme. In fact, canonical LysM/YkuD LDTs are absent in the *L. pneumophila* genome, so we investigated its potential function in maintaining cell wall integrity.

An in-frame deletion mutation was constructed (*Δlpg1697*) and tested for growth in broth culture and survival after prolonged water exposure. Growth in broth was indistinguishable for the deletion and the WT (Fig. 5A), however, after 51 days of incubation in water at 42°C, survival of the mutant was less than 10^3^ that of the WT (Fig. 5A). Furthermore, the mutant clearly lost viability relative to the WT by 14 days post-incubation. The *lpg1697* gene is located directly adjacent to the *rsmZ* noncoding RNA in *L. pneumophila*. To exclude polar effects of *lpg1697* deletion on *rsmZ* expression we analyzed expression of *lpg1697, rsmZ*, as well as the unlinked *vipD* and *rrnB* genes as controls after growth in broth (Fig. 5C). With the exception of *lpg1697*, expression of each gene in the mutant background was comparable to the WT (Fig. 5C). Transcription was restored by complementation of *lpg1697* in the chromosome (36) (Fig. 5C).

**Fig. 5.**
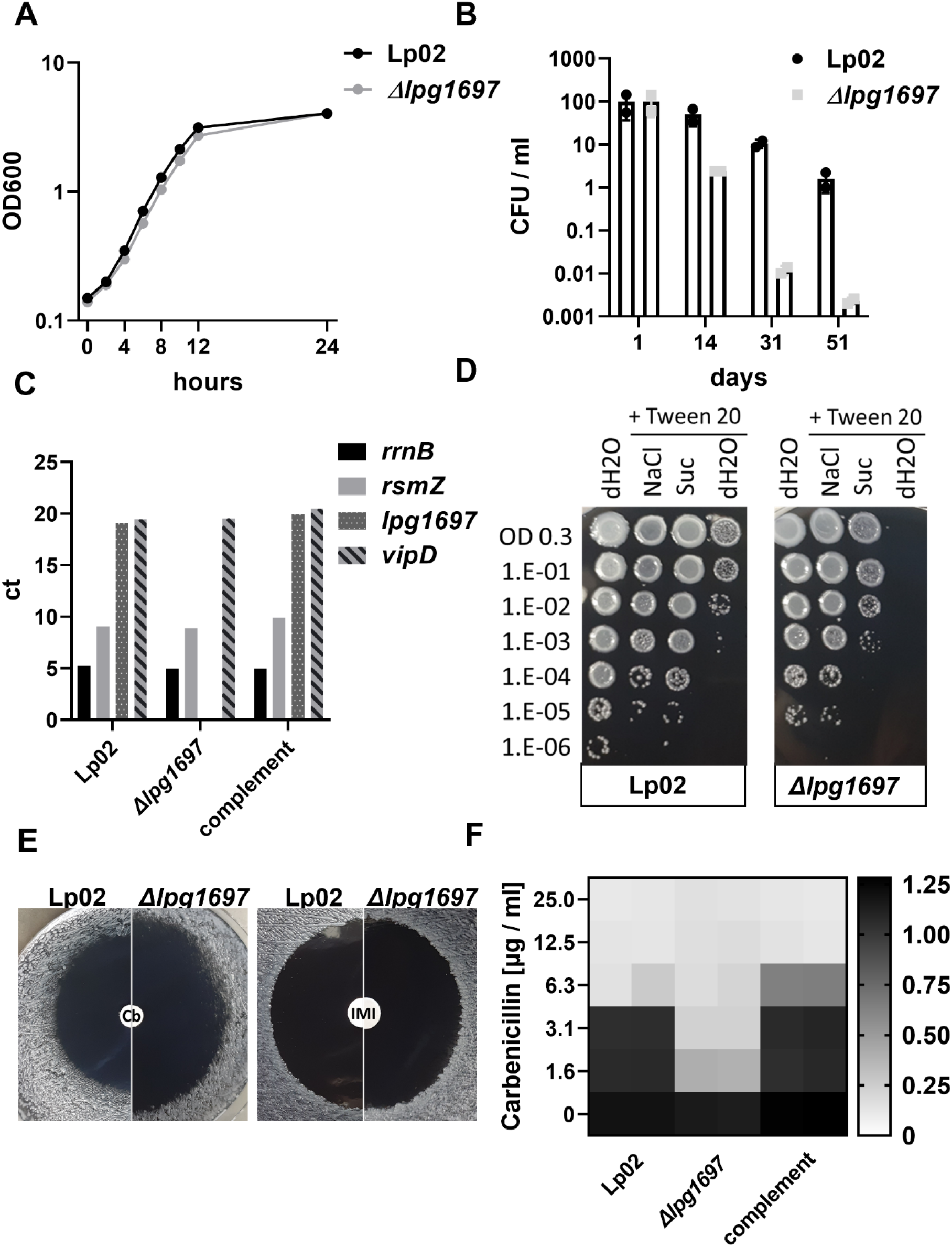
Loss of Lpg1697 results in defective survival in water as well as hypersensitivity to carbenicillin and osmotic stress. A) The *Δlpg1697* shows no fitness defect during broth growth. B) Water stress results in decreased viability of *L. pneumophila Δlpg1697* relative to WT. C) Displayed is qRT PCR analysis of four genes in the noted stains. *L. pneumophila Δlpg1697* shows no detectable transcription of *lpg1697*. Transcript levels of *rsmZ* are similar in all strains tested. Complement: *L. pneumophila Δlpg1697* carrying *lpg1697* in a neutral genomic site (36). D) *Δlpg1697* mutants are hypersensitive to Tween-20. Addition of osmostabilizers suppressed this effect. E) The *Δlpg1697* mutant is hypersensitive to carbenicillin. Disc diffusion assays were performed with carbenicillin (left) and imipenem (right), as described in Materials and Methods. F) The *Δlpg1697* mutant has lowered MIC to Cb relative to wildtype. Scale bar: A_600_ after 46 hours of growth measured in 96 well plate.

Given the presence of a predicted LDT domain in Lpg1697, we next tested deletion mutants for cell envelope phenotypes by exposure to envelope stress and to cell wall targeting antibiotics. After growth in broth to postexponential phase (PE) and 30 min treatment with 0.05 % TWEEN-20 in ultrapure water, the CFU of *Δlpg1697* strain decreased by 10^3^ compared to the WT (Fig. 5D). Addition of salt (300 mM NaCl) or sucrose (300 mM) to the 0.05 % TWEEN-20 solution suppressed the effects of the detergent/hypotonic solution. 300 mM NaCl was particularly effective, with viability restored to that of WT, consistent with loss of Lpg1697 leading to a loss of cell envelope integrity.

LDTs in other organisms are resistant to most carboxypenicillins such as Cb, but can be inhibited by carbapenems, such as Imipenem (Imp) (37–39). To further discriminate Lpg1697 function, we performed a zone of inhibition assay, comparing sensitivity to the carbapenem antibiotic Imipenem (Imp) with Cb (Fig 5E). The deletion mutant was hypersensitive to Cb, showing an increased zone of inhibition relative to the WT, whereas in the presence of Imp, there was no distinguishable difference between the mutant and WT (Fig. 5E). Concordantly, MIC Cb decreased 4-fold in *Δlpg1697*, but was comparable to the WT in the complementing strain (Fig. 5F). Therefore, loss of Lpg1697 results in a clearly definable loss in resistance to an antibiotic that shows higher efficacy against other known penicillin binding proteins, consistent with Lpg1697 having LDT activity.

We therefore next analyzed murein composition of the WT and deletion strains by ultra high-performance liquid chromatography (UHPLC) (40). PE phase Lpn was used because growth in these conditions resulted in enhanced antibiotic and osmotic-challenge hypersensitivity of the mutant (Fig. 5D-F). Surprisingly, the murein profile of *Δlpg1697* resembled the wildtype profile without detectable changes in either muropeptide patterns, the fraction of L,D-crosslinks, and overall crosslinking (Fig. S2). This is consistent with the absence of Lpg1697 causing subtle differences in peptidoglycan structure that result in envelope instability that are difficult to detect using this particular strategy.

### Loss of Lpg1697 function results in defect in intracellular growth

Having established that loss of *lpg1697* negatively impacts the capability of Lpn to persist during water stress, we were interested if its loss also reduces intracellular growth fitness after culturing to PE phase under permissive conditions. To this end, the WT and isogenic type 4 secretion system (T4SS) mutants were compared to the *Δlpg1697* strain by challenging both *A. castellanii* and murine bone marrow derived macrophages (BMDM) at MOI = 0.05 and 0.01, respectively (Fig. 6). Strikingly, intracellular growth of the *Δlpg1697* mutant was severely attenuated in amoebae with 100-fold lower CFU at 72 hpi compared to the wildtype or the strain harboring the complementing gene (Fig. 6A). There was a similar level of attenuation of intracellular growth during BMDM challenge, indicating the Lpg1697 is required for high persistence in water as well as high fitness after uptake into host cells.

**Fig. 6.**
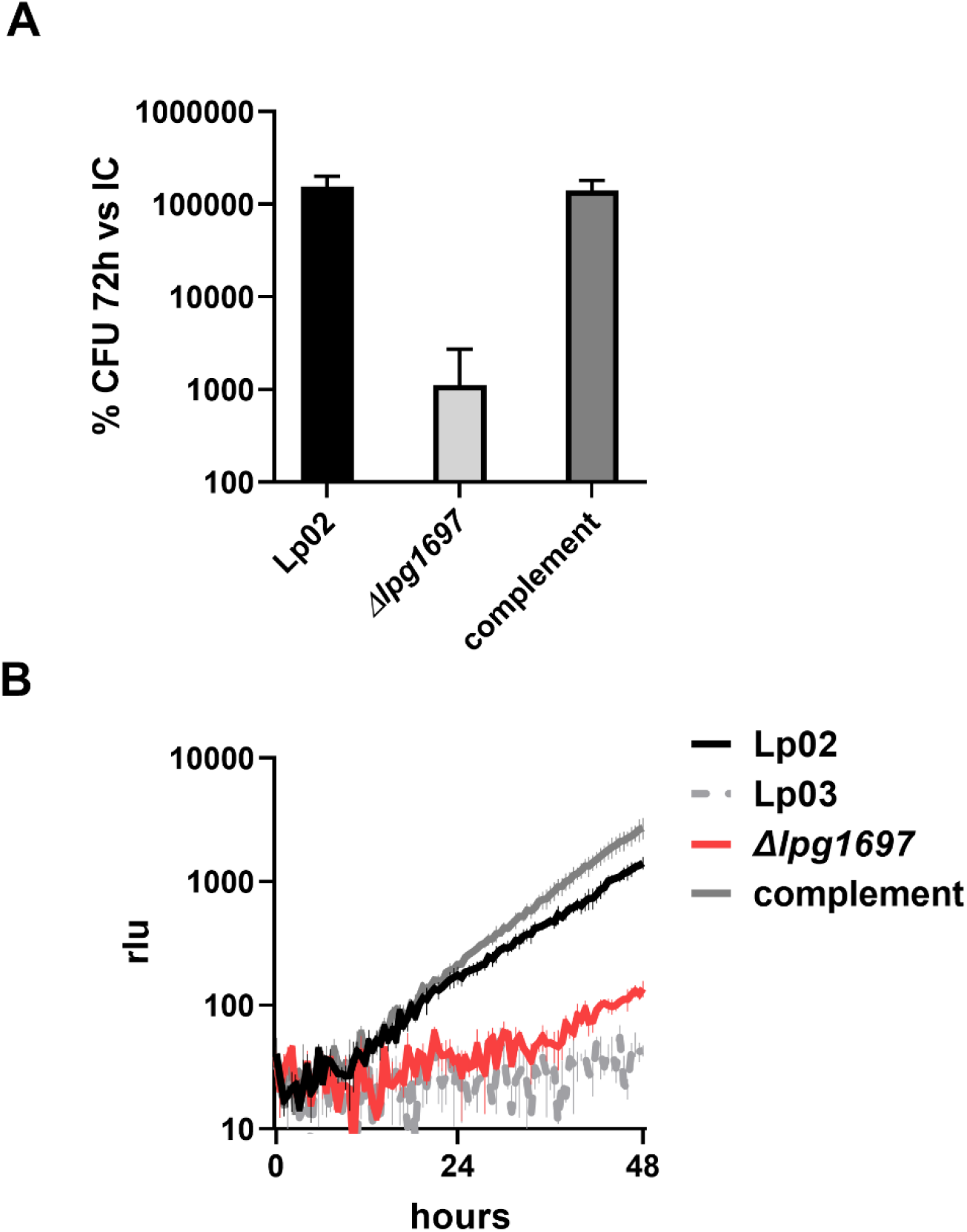
Lpg1697 function is required for efficient intracellular growth in *A. castellanii* and macrophages. A) Impaired growth of *Δlpg1697* within *A. castellanii*. Growth assayed by CFU on CYE plates 72 hr postinfection. Complement: *L. pneumophila Δlpg1697* carrying *lpg1697* in a neutral genomic site (36). B) Impaired intracellular growth of Lpn *Δlpg1697* in murine bone marrow derived macrophages and restoration of growth by the complementing strain (MOI=0.01).

Taken together, we confirmed essentiality of twelve previously unrecognized water stress-essential genes of *L. pneumophila*. Our data highlight an essential role of a YkuD domain protein for persistence and intracellular growth in *A. castellanii* and macrophages. To our knowledge this is the first report highlighting conditional essentiality of a YkuD domain protein in *Legionella* without an apparent link to peptidoglycan structure.

## Discussion

As a waterborne pathogen, *L. pneumophila* transmission to humans is tightly linked to its survival in tap water. Since *Lpn* has complex nutritional requirements including auxotrophies for at least eight amino acids (41) it cannot proliferate in the absence of appropriate nutrients or a permissive host. In spite of these restrictions, it is capable of persisting for more than a year in water (42–44). We here provide a comprehensive screen for genes that are essential for this process. Tn-seq analysis revealed 45 genes whose interruptions were not tolerated by starving *L. pneumophila*. Using CRISPR interference we showed that a large number of these genes are essential for survival in clonal populations while the remaining genes might constitute fitness factors with an essential role only under competitive conditions. Our data extend the identification of Achilles heel genes required under water stress conditions, uncovering novel vulnerabilities of the pathogen in its persistent state.

Previous studies indicated the CsrR regulator is an important persistence factor for Lpn, and a reciprocal model in which CsrR governs Lpn environmental resilience while its paralog repressor CsrA (12) directs its intracellular replication-transmission cycle has been proposed (15). In addition, an RpoS-dependent operon *lpg0279-77* encoding a two-component signaling system and a putative regulator has been reported to support survival in a nutrient-scarce environment (45). Our transposon analysis could not evaluate the importance of CsrR during water starvation, as the number of AT targets was too small to make an accurate call. On the other hand, the RpoS-regulated operon appeared nonessential in either broth or water starvation conditions (Supplemental Dataset 1). Similarly, LasM and a putative transcriptional regulator *lpg2524* both promote survival of Lpn in artificial freshwater (16, 17), but we could find no evidence for fitness defects in tap water (Supplemental Dataset 1). Other previous work concluded a role for the stringent response and RpoS, as well as the LetA/S two-component signaling systems for successful persistence in water by orchestration of a broad transcriptomic downshift concurrent with the activation of transmissive traits (14, 23, 46). Both the *rpoS* as well as the *letAS* genes were considered essential for long-term water survival based on our Tn-seq analysis, with no survival of these mutants after 50 days in water. Similarly, we found evidence for the importance of the type 2 protein secretion system (TT2S) Lsp, which was identified as contributing to fitness in cold water (10). We detected mutations in *lspI* and *lspC* reducing fitness in our Tn-seq screen performed at 42°C, indicating that the importance of T2SS might not be limited to cold water. Our work further supports a previously described role for EnhB/C in maintaining envelope integrity (47) and adaptation to freshwater (18).

Previous predictions of genes important for water survival based on analyses differential transcription in response to water stress have been partially supported by the work reported here. For instance, there is a general trend of gene downregulation after introduction into water that is countered by upregulation of *enh* genes, which we have shown are essential for viability after water starvation (18). On the other hand, 20 of the candidate essential genes identified in this work were more than 4-fold downregulated after 24 hours starvation in that study (18) (Table S3). This indicates that although essential persistence factors may have low expression levels during starvation, they are doing so in a background of general transcriptional repression. In addition, there is the paradoxical result that six genes identified as essential for water stress resistance in our work, including *ppsA* (*lpg0805*) and *mip* (*lpg0791*), were upregulated in *letS*-cells after experiencing water stress for 2h (14). This is surprising and suggests that the transcriptionally negative effect of *letS* on these genes in water may be transient (48), further arguing that the timepoint of transcriptional analysis after starvation is important for the analysis. This is in line with another study analyzing the transcriptome of RpoS mutants in water (23), in which three of our candidate genes, *ykuD/lpg1697, uvrB/lpg0072*, and Lon protease gene (*lpg1859*) are activated while two, i.e. *ppsA* and *enhC* are repressed by RpoS. Our study suggests that absence of either of these genes results in an attenuated capability to persist, so there may be temporal regulation of these genes that has been previously unappreciated.

One of the most striking results of this screen was the identification of numerous genes critical for respiratory chain function, mostly related to cytochrome C. This undoubtedly reflects the ATP-limiting conditions encountered during extended water exposure, and highlights the critical roles that maintenance of membrane potential as well as intact ATP homeostasis play to support survival in water starvation, as demonstrated in earlier studies (49, 50). Interestingly, depletion of intracellular ATP has been correlated with high drug persistence (51, 52). Concordantly, respiratory chain mutations were recently identified in screens for strains having high drug persistence, and a model linking defects in respiratory chain function to cytoplasmic acidification, biosynthetic imbalance and eventual hyperstimulation of the RpoS response has been proposed (53). Strikingly, our screen for genes whose interruption is lethal in a growth limiting condition uncovered multiple respiratory chain genes which highlights the costs associated with drug tolerance and exemplifies the selective forces favoring phenotypic variation over genetic mutation in persister cells (54).

Maintenance of cell envelope integrity is a key microbial strategy for stress adaptation (55–57). Therefore, a putative cell envelope-associated Lpg1697 that is essential for persistence in water attracted our attention. Members of this family are present in a wide range of bacteria and can act as L,D-transpeptidases (LDT) catalyzing L-D type peptidogylcan crosslinks that act as alternatives to D-D type crosslinks made by penicillin binding proteins. The catalytic YkuD domain proteins use an active site cysteine, and usually have a two-domain architecture with an N-terminal LysM peptidoglycan binding domain and a C-terminal catalytic domain. With the exception of carbapenems, most β-lactam antibiotics cannot access the active site, allowing activity in the presence of most β-lactams (58, 59). Therefore, loss of this protein is predicted to have little effect on bacterial sensitivity to carbapenems, but should result in loss of a critical barrier to protecting against carbenicillin sensitivity. Consistent with Lpg1697 have an important physiological function, hypersensitivity to carbenicillin was observed after deletion of these protein (Fig. 5) (39, 60). Enhanced sensitivity as a consequence of loss of LDT activity has been implicated in protection against envelope stress in other organisms (61–63). In Mycobacteria it was suggested that D-D linkages are likely less flexible than L-D linkages, so the interplay between these two crosslinks could be an important determinant of the stress response to hypo-osmolar conditions (64). Indeed, we observed a severe persistence defect of both the *crlpg1697* and *Δlpg1697* strains, with rapid loss of culturable bacteria after prolonged water stress (Fig. 4, 5).

Despite the clearly definable phenotypes with characteristics typical for loss of L,D-transpeptidase function, murein profiling showed no detectable compositional differences of *Δlpg1697* relative to the WT. Although this failure could be due to limitations of the technique, it should be noted that there are 10 Lpn genes encoding putative YkuD domain proteins that could compensate for loss of *lpg1697* and mask alterations in peptidoglycan profile (Fig. S2). However, given the clearly definable phenotypes resulting from loss of Lpg1697 activity, there could be an alternative function unrelated to LDT activity and not detectable by muropeptide profiling. As is true of other bacteria lacking Braun’s lipoprotein (Lpp), Legionella species employ outer membrane beta barrel proteins for cell wall anchoring of the outer membrane in a process that partially depends on LDT activity (35). It is possible that Lpg1697 is involved in crosslinks that are specific to anchoring activities constituting only a small fraction of the total murein crosslinks in the envelope. The phenotypes observed in this work could originate from a poorly tethered OM resulting from a spatially-restricted defect in crosslinking.

In conclusion, the strategy described here is the first exhaustive description of Lpn proteins essential for persistence in water that are dispensable for growth in culture. This work pointed toward the importance of the electron transport change in maintaining viability as well as the identification of a putative Ldt that likely acts to maintain cell envelope integrity. Surprisingly, this protein is also required for intracellular growth, providing a surprising link between surviving hypotonic stress and establishing an intracellular niche.

## Materials and Methods

### Bacterial strains and growth conditions

Throughout the study derivatives of *Legionella pneumophila* Philadelphia-1 strain Lp02Thy^+^ were used (28). In Lp02Thy^+^, the chromosomal *thy* mutant allele in the parental Lp02 strain was replaced with a functional *thyA* allele by allelic exchange using pJB3395 as previously described (65, 66). *L. pneumophila* strains were cultured at 37°C in BYEα broth or BCYEα agar plates (67) containing 0.4 mg/ml L - cysteine sulfate (Sigma), 0.135 mg/ml ferric nitrate (Sigma), 0.1% (w/v) α-ketoglutarate and, when appropriate, 0.1 mg/ml thymidine (Sigma), 40 μg/ml kanamycin, or 5% sucrose. Plasmids were introduced into *L. pneumophila* by electroporation (68).

*Escherichia coli* strain DH5α λpir was used for all cloning. E. coli strains were grown in Luria broth (LB) or on solid LB plates supplemented with 50 μg/ml ampicillin, 25 μg/ml chloramphenicol, or 50 μg/ml kanamycin when appropriate. Bacterial strains used in this study are summarized in Table S4. Primary bone marrow-derived macrophages from A/J mice were isolated and cultured as previously described (28). *A. castellanii* (ATCC 30234) was cultured as previously described (69).

### Construction of transposon mutant libraries

Mutagenesis of *L. pneumophila* Lp02Thy^+^ with the Mariner *himar1* transposon was performed by electroporating pTO100MmeI in triplicate as described previously (70, 71). Transformants were selected on 150mm agar plates containing kanamycin. After colonies were washed off with BYEα medium, individual pools were stored at −80°C after adding glycerol to 25% (v/v).

### Illumina library preparation and sequencing for Tn-seq

Bacterial genomic DNA was extracted with a DNeasy kit (Qiagen) and quantified by a SYBR green microtiter plate assay. Tagmentation and amplification of transposon-adjacent DNA for Illumina sequencing was conducted using a modified Nextera DNA Library Prep method as described previously (72). Samples were multiplexed, reconditioned, and size selected (250 or 275 to 600 bp; Pippin HT) before sequencing (single-end 50 bp) using custom primers (72, 73) on a HiSeq2500 with high-output V4 chemistry at the Tufts University Genomics Core Facility.

### Establishing water cultures

For the Tn-seq screen, pools of transposon mutants were directly inocculated from frozen stocks, adjusted to an A_600_=0.2, and grown overnight to an A_600_=3.8. Subsequently, cultures were washed twice in sterile tap water and inoculated to A_600_=0.03 in 200ml water in 250ml flasks. All tap water used was first passed through a 0.2μm filter (Steritop, Millipore) and then autoclaved (74, 75). 250ml flasks were autoclaved twice with tap water prior to usage. Water cultures were stored in a humidified incubator at 42°C. Before plating, flasks were carefully swirled 25 times.

For confirmation of candidate genes by CRISPRi, single colonies of *L. pneumophila* Philadelphia-1 *Lp02thyA::dcas9 PahpC::luxCDABE* containing a single CRISPRi construct were grown and processed as above with the difference that the culture volume was reduced to 20ml water held in 50ml glass tubes. For evaluation of long term efficiancy of CRISPRi with the *dotO* gene, the water culture was established in the identical fashion with the difference that ambient temperature (22°C) was used for incubation to allow for assessment of phenotype in a timeframe exceeding 50 days.

### Generation of L. pneumophila CRISPRi strains

Plasmids encoding crRNAs for targeting single ORFs by CRISPRi were constructed as described elsewhere (27). Oligos containing *Bsa*I overhangs were ordered from Integrated DNA Technologies (IDT), duplexed, and ligated into pMME1540*Bsa*I to create preliminary plasmids which were confirmed by Sanger sequencing. Final plasmids were constructed by introducing pMME985 (tracrRNA) and a preliminary plasmid into pMME977 by the Gateway LR reaction. After confirmation of single clones, the final plasmids were electroporated into *L. pneumophila* Philadelphia-1 Lp02dCas9Lux and selected on plates without thymidine (27).

### Tn-seq data analysis

Sequencing reads were processed using FASTX-toolkit and BWA (Burroughs-Wheeler Aligner), built-in aligner in TPP was used to map reads into the chromosome (AE017354). Pre-processed reads were further analyzed by TRANSIT using the Gumbel subroutine (22). Transposon insertion sites located in the 5’- and 3’-terminal 10% of the open reading frame were excluded and Gumbel was used to determine conditional gene essentiality.

### Molecular cloning and mutant construction

Gene deletions were conducted by tandem double recombination using the suicide plasmid pSR47s as described previously (76), all strains used are listed in Tab. S4. Plasmids were propagated in *E. coli* DH5α λpir (77). For complementation, the *lpg1697* gene was integrated into the intergenic region between *lpg2528* and *lpg2529* as described previously (36). All plasmids were confirmed by sequencing and are listed in Tab. S4.

### qRT-PCR gene expression analysis

Bacterial cultures were fixed in Bacterial RNA Protect reagent (Qiagen) at the timepoint of water culture inoculation and stored at −20°C until use. RNA was extracted by a RNeasy Kit (Qiagen, 74106), and cDNA was synthesized using SuperScript VILO cDNA kit (Invitrogen, 11754050). Subsequently, cDNA was amplified with PowerUp SYBR Green Master Mix (Applied Biosystems, A25742) in a StepOnePlus Real-Time PCR system (ABI, 4376600) following the manufacturer’s instructions. Primers were designed for amplification of 150-200 bp fragments using the Geneious software package and are listed in Tab. S2. Controls including absence of reverse transcription were included to confirm absence of gDNA contaminants. For normalization, *rrnB* (encoding 16S ribosomal RNA) primers were included. Transcript levels of each targeted gene were calculated by 2^-ΔΔCt^ method relative to the reference strain carrying a nontargeting control plasmid.

### Intracellular growth assays

Growth of *L. pneumophila* within *A. castellanii*, and A/J bone marrow-derived primary murine macrophages was monitored as described previously (28, 31, 71). If cocultures were plated for CFU, cells were lysed by 0.1% saponin followed by pipetting.

### Sacculi isolation and peptidoglycan analysis

Peptidoglycan (PG) analysis was performed as previously described (78). In short, *L. pneumophila* cells were centrifuged, resuspended in PBS + 5% SDS and boiled for 2 h while stirring. To remove SDS from the sacculi, samples were ultracentrifuged and washed with MilliQ water several times. After the washes, sacculi were resuspended in 100 mM Tris-HCl (pH=8) buffer with proteinase K (20 μg/ml) and incubated at 37°C for 1 hour. After that, the reaction was stopped by adding SDS (1% final concentration) and boiling at 100°C for 5 minutes. SDS was removed using described multiple washes, and sacculi were resuspended in water. Muramidase was added to the samples, which were incubated at 37°C overnight, solubilizing muropeptides from the sacculi. Soluble muropeptides were reduced by adding NaBH4 and pH was adjusted to pH=3. Muropeptides were separated using ultraperformance liquid chromatography (UPLC; Waters) and later identified using a matrix-assisted laser desorption ionization-time of flight mass spectrometry system (MALDI-TOF MS, Waters). Triplicates from the PG profiles for each representative strain were chosen for quantification, and the area under each peak was integrated for statistical analysis.

### Osmotic tolerance studies

Osmotic challenge assays were performed as described previously with modifications (79). Briefly, fresh postexponential phase cultures (OD600 3.5) were adjusted to an OD600 = 0.3, and 1 ml suspensions were pelleted by centrifugation at low speed (5 min 5000 × g). Subsequently, supernatants were removed and replaced with the indicated stress solutions and distilled water as a control. Cells were incubated in these conditions for 30 min in a shaking incubator at 37°C. Viability was determined by spot plating dilution rows on BCYEα agar plates.

### Antimicrobial susceptibility testing

Disc diffusion assays were performed as described previosly with modifications (80). Briefly, 6mm filter discs were impregnated with 5 μl of a freshly prepared imipenem solution (2mg / ml in H2O) or 5 μl of a 50 mg / ml carbenicillin solution, respectively. The discs were then placed on fresh BCYEα agar plates containing a fresh layer of the indicated *Legionella* strain. After three days of incubation, zones of inhibition were determined.

MIC carbenicillin was determined by microtiter plate assays as described previously with modifications (81). Briefly, two single colonies of each *L. pneumophila* strain were pregrown to exponential growth phase in BYEα broth and then back-diluted to an OD600 of 0.15. 100μl aliquots of this suspension were distributed into a 96-well plate preloaded with a concentration gradient of serial 2-fold carbenicillin dilutions in BYEα (100 μl / well). The plates were then grown for 46 hours in a plate reader with continuous shaking (Tecan Infinite 200 PRO) and bacterial growth was recorded as optical density. MICs were determined as the lowest antibiotic concentration for which no growth of *L. pneumophila* was detected after 46 h at 37°C. Wells containing BYEα broth or bacterial suspension without antibiotic were used as negative and positive growth controls, respectively.

## Funding Acknowledgements

This work was supported by the Deutsche Forschungsgemeinschaft (DFG, German Research Foundation) grant AU550/1-1 to PA and NIAID award R01 AI146245 to RRI. Research in the Cava lab is supported by the Knut and Alice Wallenberg Foundation (KAW), The Laboratory of Molecular Infection Medicine Sweden (MIMS), the Swedish Research Council and the Kempe Foundation.

## Acknowledgements

We thank the M. Machner lab for providing *L. pn* MML109, and the pMME plasmids for CRISPRi.

## Data availability

The data for this study have been deposited in the NCBI sequence read archive under accession number PRJNA816157 (https://www.ncbi.xxx).

## Figure Legends

**Fig. S1.**
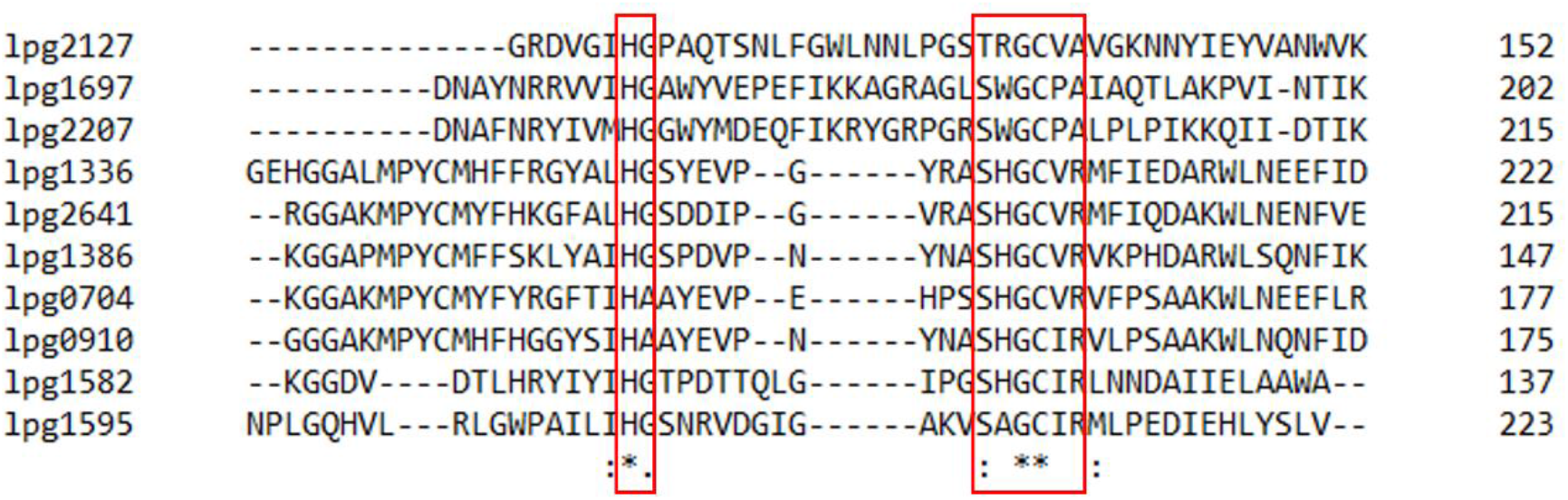
Partial ClustalOmega amino acid sequence alignment of YkuD like domain proteins of *Lpn*. Conserved residues involved in catalysis are boxed red.

**Fig. S2.**
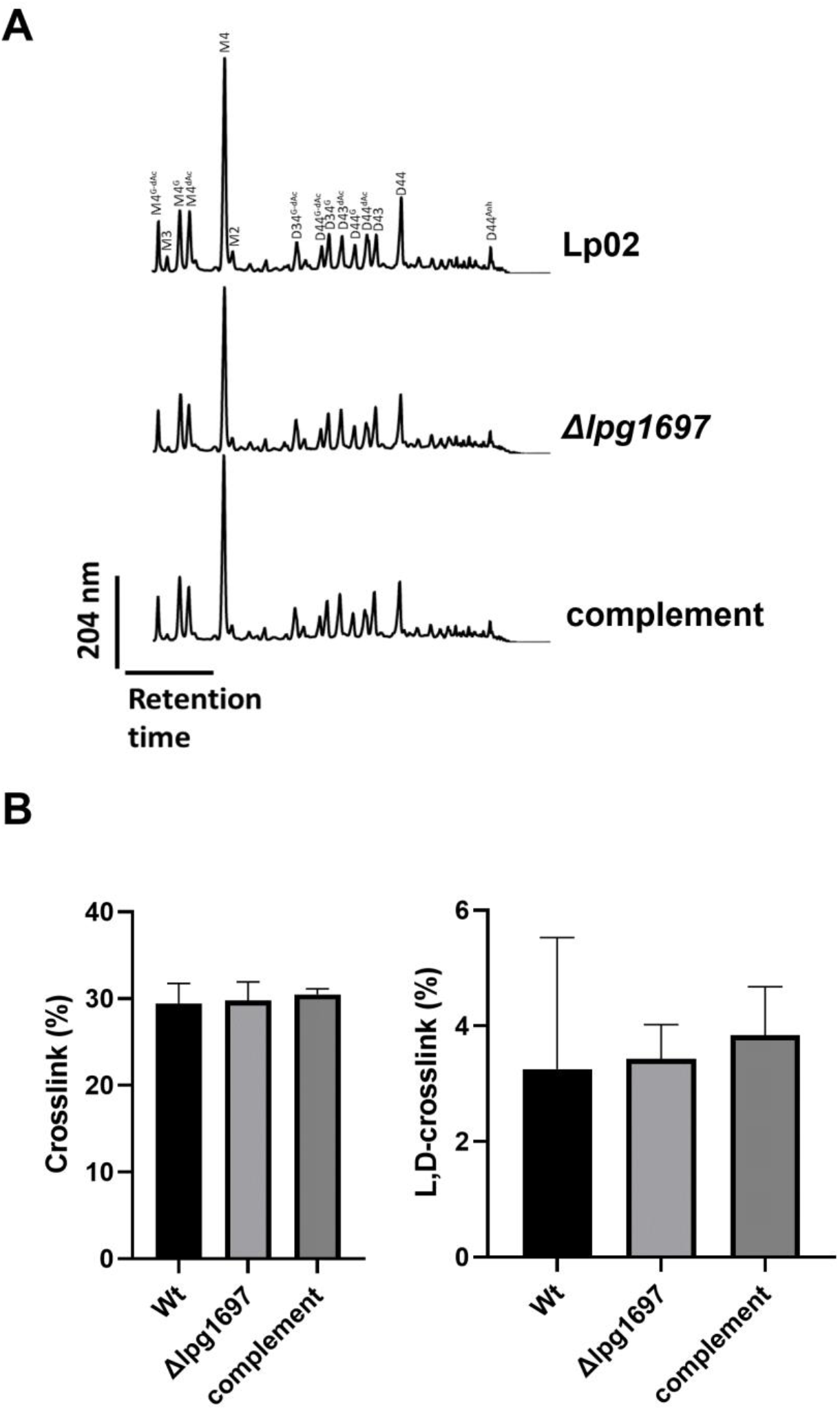
Absence of detectable changes in muropeptide profiles related to loss of *lpg1697* gene. A) chromatograms of UHPLC runs B) relative amount of total crosslinkages and L,D-crosslinkages were not significantly different between the strains (two tailed unpaired t-test)

## References

1. Collier SA, Deng L, Adam EA, Benedict KM, Beshearse EM, Blackstock AJ, Bruce BB, Derado G, Edens C, Fullerton KE, Gargano JW, Geissler AL, Hall AJ, Havelaar AH, Hill VR, Hoekstra RM, Reddy SC, Scallan E, Stokes EK, Yoder JS, Beach MJ. 2021. Estimate of Burden and Direct Healthcare Cost of Infectious Waterborne Disease in the United States. Emerg Infect Dis 27:140–149.

2. Fields BS, Benson RF, Besser RE. 2002. Legionella and Legionnaires’ disease: 25 years of investigation. Clin Microbiol Rev 15:506–26.

3. Hilbi H, Hoffmann C, Harrison CF. 2011. Legionella spp. outdoors: colonization, communication and persistence. Environ Microbiol Rep 3:286–96.

4. Falkinham JO, 3rd. 2020. Living with Legionella and Other Waterborne Pathogens. Microorganisms 8.

5. Zhang C, Struewing I, Mistry JH, Wahman DG, Pressman J, Lu J. 2021. Legionella and other opportunistic pathogens in full-scale chloraminated municipal drinking water distribution systems. Water Res 205:117571.

6. https://www.cdc.gov/legionella/health-depts/surv-reporting/surveillance-reports.html.

7. Schofield GM. 1985. A note on the survival of Legionella pneumophila in stagnant tap water. J Appl Bacteriol 59:333–5.

8. Paszko-Kolva C, Shahamat M, Colwell RR. 1993. Effect of temperature on survival of Legionella pneumophila in the aquatic environment. Microb Releases 2:73–9.

9. Ohno A, Kato N, Yamada K, Yamaguchi K. 2003. Factors influencing survival of Legionella pneumophila serotype 1 in hot spring water and tap water. Appl Environ Microbiol 69:2540–7.

10. Soderberg MA, Dao J, Starkenburg SR, Cianciotto NP. 2008. Importance of type II secretion for survival of Legionella pneumophila in tap water and in amoebae at low temperatures. Appl Environ Microbiol 74:5583–8.

11. Hammer BK, Tateda ES, Swanson MS. 2002. A two-component regulator induces the transmission phenotype of stationary-phase Legionella pneumophila. Mol Microbiol 44:107–18.

12. Molofsky AB, Swanson MS. 2003. Legionella pneumophila CsrA is a pivotal repressor of transmission traits and activator of replication. Mol Microbiol 50:445–61.

13. Sahr T, Rusniok C, Dervins-Ravault D, Sismeiro O, Coppee JY, Buchrieser C. 2012. Deep sequencing defines the transcriptional map of L. pneumophila and identifies growth phase-dependent regulated ncRNAs implicated in virulence. RNA Biol 9:503–19.

14. Mendis N, McBride P, Saoud J, Mani T, Faucher SP. 2018. The LetA/S two-component system regulates transcriptomic changes that are essential for the culturability of Legionella pneumophila in water. Sci Rep 8:6764.

15. Abbott ZD, Yakhnin H, Babitzke P, Swanson MS. 2015. csrR, a Paralog and Direct Target of CsrA, Promotes Legionella pneumophila Resilience in Water. mBio 6:e00595.

16. Li L, Faucher SP. 2016. The Membrane Protein LasM Promotes the Culturability of Legionella pneumophila in Water. Front Cell Infect Microbiol 6:113.

17. Li L, Faucher SP. 2017. Role of the LuxR family transcriptional regulator Lpg2524 in the survival of Legionella pneumophila in water. Can J Microbiol 63:535–545.

18. Li L, Mendis N, Trigui H, Faucher SP. 2015. Transcriptomic changes of Legionella pneumophila in water. BMC Genomics 16:637.

19. Aurass P, Gerlach T, Becher D, Voigt B, Karste S, Bernhardt J, Riedel K, Hecker M, Flieger A. 2016. Life Stage-specific Proteomes of Legionella pneumophila Reveal a Highly Differential Abundance of Virulence-associated Dot/Icm effectors. Mol Cell Proteomics 15:177–200.

20. Swanson MS, Hammer BK. 2000. Legionella pneumophila pathogesesis: a fateful journey from amoebae to macrophages. Annu Rev Microbiol 54:567–613.

21. Mondino S, Schmidt S, Rolando M, Escoll P, Gomez-Valero L, Buchrieser C. 2020. Legionnaires’ Disease: State of the Art Knowledge of Pathogenesis Mechanisms of Legionella. Annu Rev Pathol 15:439–466.

22. DeJesus MA, Ambadipudi C, Baker R, Sassetti C, loerger TR. 2015. TRANSIT--A Software Tool for Himar1 TnSeq Analysis. PLoS Comput Biol 11:e1004401.

23. Trigui H, Dudyk P, Oh J, Hong JI, Faucher SP. 2015. A regulatory feedback loop between RpoS and SpoT supports the survival of Legionella pneumophila in water. Appl Environ Microbiol 81:918–28.

24. Lampe DJ, Grant TE, Robertson HM. 1998. Factors affecting transposition of the Himar1 mariner transposon in vitro. Genetics 149:179–87.

25. Chien M, Morozova I, Shi S, Sheng H, Chen J, Gomez SM, Asamani G, Hill K, Nuara J, Feder M, Rineer J, Greenberg JJ, Steshenko V, Park SH, Zhao B, Teplitskaya E, Edwards JR, Pampou S, Georghiou A, Chou IC, Iannuccilli W, Ulz ME, Kim DH, Geringer-Sameth A, Goldsberry C, Morozov P, Fischer SG, Segal G, Qu X, Rzhetsky A, Zhang P, Cayanis E, De Jong PJ, Ju J, Kalachikov S, Shuman HA, Russo JJ. 2004. The genomic sequence of the accidental pathogen Legionella pneumophila. Science 305:1966–8.

26. Al-Bana BH, Haddad MT, Garduno RA. 2014. Stationary phase and mature infectious forms of Legionella pneumophila produce distinct viable but non-culturable cells. Environ Microbiol 16:382–95.

27. Ellis NA, Kim B, Tung J, Machner MP. 2021. A multiplex CRISPR interference tool for virulence gene interrogation in Legionella pneumophila. Commun Biol 4:157.

28. Berger KH, Isberg RR. 1993. Two distinct defects in intracellular growth complemented by a single genetic locus in Legionella pneumophila. Mol Microbiol 7:7–19.

29. Bergkessel M. 2021. Bacterial transcription during growth arrest. Transcription 12:232–249.

30. Bai J, Dai Y, Farinha A, Tang AY, Syal S, Vargas-Cuebas G, van Opijnen T, Isberg RR, Geisinger E. 2021. Essential Gene Analysis in Acinetobacter baumannii by High-Density Transposon Mutagenesis and CRISPR Interference. J Bacteriol 203:e0056520.

31. Coers J, Vance RE, Fontana MF, Dietrich WF. 2007. Restriction of Legionella pneumophila growth in macrophages requires the concerted action of cytokine and Naip5/Ipaf signalling pathways. Cell Microbiol 9:2344–57.

32. Ensminger AW, Yassin Y, Miron A, Isberg RR. 2012. Experimental evolution of Legionella pneumophila in mouse macrophages leads to strains with altered determinants of environmental survival. PLoS Pathog 8:e1002731.

33. Falls KC, Williams AL, Bryksin AV, Matsumura I. 2014. Escherichia coli deletion mutants illuminate trade-offs between growth rate and flux through a foreign anabolic pathway. PLoS One 9:e88159.

34. Amano K, Williams JC. 1983. Peptidoglycan of Legionella pneumophila: apparent resistance to lysozyme hydrolysis correlates with a high degree of peptide cross-linking. J Bacteriol 153:520–6.

35. Sandoz KM, Moore RA, Beare PA, Patel AV, Smith RE, Bern M, Hwang H, Cooper CJ, Priola SA, Parks JM, Gumbart JC, Mesnage S, Heinzen RA. 2021. beta-Barrel proteins tether the outer membrane in many Gram-negative bacteria. Nat Microbiol 6:19–26.

36. Liu Y, Gao P, Banga S, Luo ZQ. 2008. An in vivo gene deletion system for determining temporal requirement of bacterial virulence factors. Proc Natl Acad Sci U S A 105:9385–90.

37. Hugonnet JE, Mengin-Lecreulx D, Monton A, den Blaauwen T, Carbonnelle E, Veckerle C, Brun YV, van Nieuwenhze M, Bouchier C, Tu K, Rice LB, Arthur M. 2016. Factors essential for L,D-transpeptidase-mediated peptidoglycan cross-linking and beta-lactam resistance in Escherichia coli. Elife 5.

38. Mainardi JL, Hugonnet JE, Rusconi F, Fourgeaud M, Dubost L, Moumi AN, Delfosse V, Mayer C, Gutmann L, Rice LB, Arthur M. 2007. Unexpected inhibition of peptidoglycan LD-transpeptidase from Enterococcus faecium by the beta-lactam imipenem. J Biol Chem 282:30414–22.

39. Aliashkevich A, Cava F. 2021. LD-transpeptidases: the great unknown among the peptidoglycan cross-linkers. FEBS J doi:10.1111/febs.16066.

40. Alvarez L, Cordier B, Van Teeffelen S, Cava F. 2020. Analysis of Gram-negative Bacteria Peptidoglycan by Ultra-performance Liquid Chromatography. Bio Protoc 10:e3780.

41. Price CT, Richards AM, Von Dwingelo JE, Samara HA, Abu Kwaik Y. 2014. Amoeba host-Legionella synchronization of amino acid auxotrophy and its role in bacterial adaptation and pathogenic evolution. Environ Microbiol 16:350–8.

42. Paszko-Kolva C, Shahamat M, Yamamoto H, Sawyer T, Vives-Rego J, Colwell RR. 1991. Survival ofLegionella pneumophila in the aquatic environment. Microb Ecol 22:75–83.

43. Paszko-Kolva C, Shahamat M, Colwell RR. 1992. Long-term survival of Legionella pneumophila serogroup 1 under low-nutrient conditions and associated morphological changes. FEMS Microbiology Ecology 11:45–55.

44. Mendis N, McBride P, Faucher SP. 2015. Short-Term and Long-Term Survival and Virulence of Legionella pneumophila in the Defined Freshwater Medium Fraquil. PLoS One 10:e0139277.

45. Hughes ED, Byrne BG, Swanson MS. 2019. A Two-Component System That Modulates Cyclic di-GMP Metabolism Promotes Legionella pneumophila Differentiation and Viability in Low-Nutrient Conditions. J Bacteriol 201.

46. Bachman MA, Swanson MS. 2004. Genetic evidence that Legionella pneumophila RpoS modulates expression of the transmission phenotype in both the exponential phase and the stationary phase. Infect Immun 72:2468–76.

47. Liu M, Conover GM, Isberg RR. 2008. Legionella pneumophila EnhC is required for efficient replication in tumour necrosis factor alpha-stimulated macrophages. Cell Microbiol 10:1906–23.

48. Cianciotto NP, Eisenstein BI, Mody CH, Engleberg NC. 1990. A mutation in the mip gene results in an attenuation of Legionella pneumophila virulence. J Infect Dis 162:121–6.

49. Rao SP, Alonso S, Rand L, Dick T, Pethe K. 2008. The protonmotive force is required for maintaining ATP homeostasis and viability of hypoxic, nonreplicating Mycobacterium tuberculosis. Proc Natl Acad Sci U S A 105:11945–50.

50. Gengenbacher M, Rao SPS, Pethe K, Dick T. 2010. Nutrient-starved, non-replicating Mycobacterium tuberculosis requires respiration, ATP synthase and isocitrate lyase for maintenance of ATP homeostasis and viability. Microbiology (Reading) 156:81–87.

51. Conlon BP, Rowe SE, Gandt AB, Nuxoll AS, Donegan NP, Zalis EA, Clair G, Adkins JN, Cheung AL, Lewis K. 2016. Persister formation in Staphylococcus aureus is associated with ATP depletion. Nat Microbiol 1:16051.

52. Shan Y, Brown Gandt A, Rowe SE, Deisinger JP, Conlon BP, Lewis K. 2017. ATP-Dependent Persister Formation in Escherichia coli. mBio 8.

53. Van den Bergh B, Schramke H, Michiels JE, Kimkes TEP, Radzikowski JL, Schimpf J, Vedelaar SR, Burschel S, Dewachter L, Loncar N, Schmidt A, Meijer T, Fauvart M, Friedrich T, Michiels J, Heinemann M. 2022. Mutations in respiratory complex I promote antibiotic persistence through alterations in intracellular acidity and protein synthesis. Nat Commun 13:546.

54. Lewis K. 2010. Persister cells. Annu Rev Microbiol 64:357–72.

55. Mitchell AM, Silhavy TJ. 2019. Envelope stress responses: balancing damage repair and toxicity. Nat Rev Microbiol 17:417–428.

56. Hews CL, Cho T, Rowley G, Raivio TL. 2019. Maintaining Integrity Under Stress: Envelope Stress Response Regulation of Pathogenesis in Gram-Negative Bacteria. Front Cell Infect Microbiol 9:313.

57. Stankeviciute G, Miguel AV, Radkov A, Chou S, Huang KC, Klein EA. 2019. Differential modes of crosslinking establish spatially distinct regions of peptidoglycan in Caulobacter crescentus. Mol Microbiol 111:995–1008.

58. Caveney NA, Caballero G, Voedts H, Niciforovic A, Worrall LJ, Vuckovic M, Fonvielle M, Hugonnet JE, Arthur M, Strynadka NCJ. 2019. Structural insight into YcbB-mediated beta-lactam resistance in Escherichia coli. Nat Commun 10:1849.

59. Dubee V, Triboulet S, Mainardi JL, Etheve-Quelquejeu M, Gutmann L, Marie A, Dubost L, Hugonnet JE, Arthur M. 2012. Inactivation of Mycobacterium tuberculosis l,d-transpeptidase LdtMt(1) by carbapenems and cephalosporins. Antimicrob Agents Chemother 56:4189–95.

60. Cordillot M, Dubee V, Triboulet S, Dubost L, Marie A, Hugonnet JE, Arthur M, Mainardi JL. 2013. In vitro cross-linking of Mycobacterium tuberculosis peptidoglycan by L,D-transpeptidases and inactivation of these enzymes by carbapenems. Antimicrob Agents Chemother 57:5940–5.

61. More N, Martorana AM, Biboy J, Otten C, Winkle M, Serrano CKG, Monton Silva A, Atkinson L, Yau H, Breukink E, den Blaauwen T, Vollmer W, Polissi A. 2019. Peptidoglycan Remodeling Enables Escherichia coli To Survive Severe Outer Membrane Assembly Defect. mBio 10.

62. Pagliai FA, Gardner CL, Bojilova L, Sarnegrim A, Tamayo C, Potts AH, Teplitski M, Folimonova SY, Gonzalez CF, Lorca GL. 2014. The transcriptional activator LdtR from ‘Candidatus Liberibacter asiaticus’ mediates osmotic stress tolerance. PLoS Pathog 10:e1004101.

63. Bernal-Cabas M, Ayala JA, Raivio TL. 2015. The Cpx envelope stress response modifies peptidoglycan cross-linking via the L,D-transpeptidase LdtD and the novel protein YgaU. J Bacteriol 197:603–14.

64. Sanders AN, Wright LF, Pavelka MS. 2014. Genetic characterization of mycobacterial L,D-transpeptidases. Microbiology (Reading) 160:1795–1806.

65. Kessler B, de Lorenzo V, Timmis KN. 1992. A general system to integrate lacZ fusions into the chromosomes of gram-negative eubacteria: regulation of the Pm promoter of the TOL plasmid studied with all controlling elements in monocopy. Mol Gen Genet 233:293–301.

66. O’Connor TJ, Zheng H, VanRheenen SM, Ghosh S, Cianciotto NP, Isberg RR. 2016. Iron Limitation Triggers Early Egress by the Intracellular Bacterial Pathogen Legionella pneumophila. Infect Immun 84:2185–2197.

67. Edelstein PH. 1981. Improved semiselective medium for isolation of Legionella pneumophila from contaminated clinical and environmental specimens. J Clin Microbiol 14:298–303.

68. Berger KH, Merriam JJ, Isberg RR. 1994. Altered intracellular targeting properties associated with mutations in the Legionella pneumophila dotA gene. Mol Microbiol 14:809–22.

69. Moffat JF, Tompkins LS. 1992. A quantitative model of intracellular growth of Legionella pneumophila in Acanthamoeba castellanii. Infect Immun 60:296–301.

70. O’Connor TJ, Adepoju Y, Boyd D, Isberg RR. 2011. Minimization of the Legionella pneumophila genome reveals chromosomal regions involved in host range expansion. Proc Natl Acad Sci U S A 108:14733–40.

71. Park JM, Ghosh S, O’Connor TJ. 2020. Combinatorial selection in amoebal hosts drives the evolution of the human pathogen Legionella pneumophila. Nat Microbiol 5:599–609.

72. Geisinger E, Vargas-Cuebas G, Mortman NJ, Syal S, Dai Y, Wainwright EL, Lazinski D, Wood S, Zhu Z, Anthony J, van Opijnen T, Isberg RR. 2019. The Landscape of Phenotypic and Transcriptional Responses to Ciprofloxacin in Acinetobacter baumannii: Acquired Resistance Alleles Modulate Drug-Induced SOS Response and Prophage Replication. mBio 10.

73. Geisinger E, Mortman NJ, Dai Y, Cokol M, Syal S, Farinha A, Fisher DG, Tang AY, Lazinski DW, Wood S, Anthony J, van Opijnen T, Isberg RR. 2020. Antibiotic susceptibility signatures identify potential antimicrobial targets in the Acinetobacter baumannii cell envelope. Nat Commun 11:4522.

74. Kampfer P, Aurass P, Karste S, Flieger A, Glaeser SP. 2016. Paracoccus contaminans sp. nov., isolated from a contaminated water microcosm. Int J Syst Evol Microbiol 66:5101–5105.

75. Aurass P, Flieger A. 2020. Complete Genome Sequence of Rhodoferax sp. Strain BAB1, Isolated after Filter Sterilization of Tap Water. Microbiol Resour Announc 9.

76. Merriam JJ, Mathur R, Maxfield-Boumil R, Isberg RR. 1997. Analysis of the Legionella pneumophila fliI gene: intracellular growth of a defined mutant defective for flagellum biosynthesis. Infect Immun 65:2497–501.

77. Kolter R, Inuzuka M, Helinski DR. 1978. Trans-complementation-dependent replication of a low molecular weight origin fragment from plasmid R6K. Cell 15:1199–208.

78. Alvarez L, Hernandez SB, de Pedro MA, Cava F. 2016. Ultra-Sensitive, High-Resolution Liquid Chromatography Methods for the High-Throughput Quantitative Analysis of Bacterial Cell Wall Chemistry and Structure. Methods Mol Biol 1440:11–27.

79. Brammer Basta LA, Ghosh A, Pan Y, Jakoncic J, Lloyd EP, Townsend CA, Lamichhane G, Bianchet MA. 2015. Loss of a Functionally and Structurally Distinct ld-Transpeptidase, LdtMt5, Compromises Cell Wall Integrity in Mycobacterium tuberculosis. J Biol Chem 290:25670–85.

80. Bauer AW, Kirby WM, Sherris JC, Turck M. 1966. Antibiotic susceptibility testing by a standardized single disk method. Tech Bull Regist Med Technol 36:49–52.

81. Vandewalle-Capo M, Massip C, Descours G, Charavit J, Chastang J, Billy PA, Boisset S, Lina G, Gilbert C, Maurin M, Jarraud S, Ginevra C. 2017. Minimum inhibitory concentration (MIC) distribution among wild-type strains of Legionella pneumophila identifies a subpopulation with reduced susceptibility to macrolides owing to efflux pump genes. Int J Antimicrob Agents 50:684–689.

